# Bioplastic biodegradability shapes microbial communities in a coastal brackish environment

**DOI:** 10.1101/2025.10.07.679016

**Authors:** Igor S. Pessi, Eeva Eronen-Rasimus, Pinja Näkki, David N. Thomas, Hermanni Kaartokallio

**Affiliations:** Marine and Freshwater Unit, Finnish Environment Institute (Syke), Helsinki, Finland; Department of Microbiology, University of Helsinki, Helsinki, Finland; Helsinki Institute of Sustainability Science (HELSUS), Helsinki, Finland; Department of Environmental Sciences, University of Helsinki, Helsinki, Finland

## Abstract

Microorganisms are metabolically versatile and central to marine ecosystems, yet the potential of marine microbial communities to degrade different bioplastics and the effect of environmental factors are poorly understood. Employing multi-seasonal *in situ* and *in vitro* experiments, we assessed the biodegradation of six commonly used bio-based bioplastic materials at a coastal site in the brackish Baltic Sea and characterised the associated microbial communities using metagenomics and metatranscriptomics. Cellulose acetate (CA), polybutylene succinate (PBS), and polyhydroxybutyrate/valerate (PHB) degraded at varying rates across materials, seasons, and experimental settings, with up to 28% weight attrition after 97 weeks *in situ* (CA) and 56% carbon loss to CO_2_ after four weeks *in vitro* (PBS). The three biodegraded plastics developed similar microbial communities that differed markedly from those on the other materials (cellulose acetate propionate, polyamide, and polyethylene) and in the water column. The main microbial populations on the biodegraded plastics included aerobic and facultative anaerobic heterotrophs with a broad capacity for carbohydrate metabolism. Populations with the potential for nitrogen fixation and denitrification were more prevalent on the biodegraded plastics, suggesting a link with the marine nitrogen cycle. Based on the metatranscriptomic signal of key genes involved in the initial hydrolysis of CA, PBS, and PHB, we identified diverse microbial populations that can potentially drive the biodegradation of these materials in the Baltic Sea, many of which encoded the potential to degrade multiple bioplastics. We propose the term *bioplastisphere* to denote the distinctive microbial communities associated with biodegradable plastics.

## Introduction

Millions of tonnes of plastics enter the oceans annually, making plastic pollution one of the most pervasive environmental challenges of our time [1–3]. Plastics accumulate in coastal regions, surface waters, and sediments, where they can cause long-lasting environmental harm [4–6]. The amount of mismanaged plastic waste in the environment is projected to triple in the coming decades due to increasing industrial and societal demands combined with inadequate waste management practices [7]. Conventional fossil fuel-based plastics, particularly polyethylene, dominate the plastic industry due to their high durability against biotic and abiotic processes, which also makes them the main constituents of marine plastic pollution [7, 8]. Bioplastics—*i*.*e*. plastics that are biodegradable or produced from renewable, bio-based sources—have become increasingly popular as part of a wider strategy to reduce environmental pollution and the use of finite fossil fuel resources [9]. However, many bioplastic materials only achieve full biodegradation under tightly constrained conditions, such as those certified by the EU standard EN13432 for industrial composting [10]. Plastic biodegradation is much slower in marine environments, which are generally colder, more oligotrophic, and contain lower microbial biomass and diversity than compost and soil [11]. Because both material properties and environmental conditions influence plastic biodegradation, there is little consensus about the general biodegradability of different bioplastic materials [12, 13]. Current industrial directives do not address adequately the impacts of mismanaged bioplastic waste in marine environments, which remain poorly understood.

Plastics can serve as substrate for microbial growth and biofilm formation in marine environments, and microorganisms are the main drivers of plastic biodegradation [14–16]. Microorganisms colonise marine plastics readily and communities are considered mature in the order of one month [17–20]. Communities associated with plastic debris, which are collectively known as the plastisphere, are generally dominated by *Alpha*- and *Gammaproteobacteria* and include potentially hydrocarbonoclastic genera [21–24]. In certain biodegradable plastics, the abundance of *Alpha*- and *Gammaproteobacteria* has been associated with the loss of substrate or observed biodegradation [18, 25, 26]. Plastisphere communities are mostly shaped by biogeographical and environmental factors [21, 22, 27– 29], and the importance of material properties is under debate [19, 22, 23, 29, 30]. Communities growing on conventional plastics are usually similar to those on totally inert surfaces (*e*.*g*. glass) and are characterised by a secondary biofilm community that is not in close contact with the substrate, with differences in community composition mostly driven by rare taxa [19, 22]. Conversely, increasing evidence suggests that microbial communities colonising some bioplastic materials (*e*.*g*. polyhydroxybutyrate and cellulose acetate) are different and more active than those on conventional plastics, whereas communities on bioplastics that have limited biodegradation in marine environments (*e*.*g*. polylactic acid) resemble those on conventional plastics [18, 24–26, 31, 32]. Recent studies have shown that metagenomics is a powerful tool to investigate the functional potential of microbial communities and their role not only in plastic biodegradation but also other biogeochemical processes. For example, increased denitrification potential as communities mature during the first months of plastic colonization has been observed in a coastal estuary [24]. This study also showed that genes for nitrogen metabolism are typically present in many common plastisphere taxa such as *Alpha*- (*e*.*g. Rhizobiales* and *Rhodobacterales*) and *Gammaproteobacteria* (*e*.*g. Burkholderiales* and *Pseudomonadales*).

Key to a better understanding of the impact of marine plastics on marine environments is to consider biodegradation as a systemic capacity, *i*.*e*. by investigating the associated microbial communities and the environmental conditions constraining their activity. However, long-term *in situ* investigations are rare, particularly studies of bioplastic biodegradation coupled with the analysis of microbial communities [19, 26, 32–37]. Data on community composition and activity at the functional level are also scarce, including not only the microbial pathways of bioplastic biodegradation but also their connections with the marine carbon and nitrogen cycles. In this study we assessed the biodegradation of six commonly used bio-based bioplastic materials at a coastal site in the brackish Baltic Sea under multi-seasonal *in situ* and *in vitro* conditions. Coupling measurements of plastic weight attrition, spectroscopy, respirometry, metagenomics, and metatranscriptomics, we investigated differences in biodegradation between the materials throughout different seasons and explored links between biodegradation and microbial community structure and activity. We hypothesised that the composition of mature (> 1 month) communities—including the presence and activity of plastic-degrading genes—is structured by the physicochemical properties of the bioplastics and therefore unique to each material.

## Materials and Methods

### Experimental setup

#### Bioplastic materials

We investigated six types of bio-based bioplastics, which will be designated as bioplastics: cellulose acetate (CA), cellulose acetate propionate (CAP), polybutylene succinate (PBS), polyhydroxybutyrate/valerate (PHB), polyamide (PA), and polyethylene (PE). The materials were selected to represent a range of commonly used bioplastics with distinct molecular structures and included *i*) natural polymers with known marine biodegradability (CA and PHB); *ii*) bio-based polymers with unclear marine biodegradability (PBS, CAP, and PA); and *iii*) non-biodegradable polymers (PE). Injection-moulded bioplastic sheets were provided by the VTT Technical Research Centre of Finland (Espoo, Finland).

#### *In situ* mesocosm

*In situ* biodegradation was investigated for 97 weeks in a mesocosm experiment set up at the Tvärminne Zoological Station, University of Helsinki (Hanko, Finland, 59.845°N, 23.250°E). The salinity of the seawater at the study site is between 5 and 6 and the water temperature during the experiment ranged from 2.5°C to 25.0ºC. In July 2021, 60 bioplastic sheets (15 mm width × 100 mm length × 3 mm thick) of each material were weighed using an EP-225-SM-DR semi-micro balance (Precisa Gravimetrics, Dietikon, Switzerland), secured to racks, and transferred to a 400 l flow-through tank connected to the Baltic Sea. Glass sheets of identical dimensions were included as control. The mesocosm was kept in the dark to prevent the growth of photosynthetic microeukaryotes. At six sampling times (September 2021, January, March, and August 2022, and January and May 2023), five sheets of each material were sampled for measuring biodegradation and five for metagenomics and metatranscriptomics. Five seawater samples (1000 ml) were also taken from inside the mesocosm tank for the molecular characterization of the planktonic communities. The samples for measuring biodegradation were kept at –20°C until processing and the other samples were processed immediately.

#### *In vitro* experiments

For the seven sampling times indicated above (July 2021–May 2023) and each bioplastic material (except CAP, which was not included due to equipment availability), 0.25 g l^–1^ of finely milled bioplastic (< 0.3 mm particle size) was added to three 600 ml amber-coloured bottles containing 250 ml of seawater sampled from the mesocosm tank. To prevent nutrient limitation, 0.1 g l^–1^ KH_2_PO_4_ and 0.5 g l^–1^ NH_4_Cl were added. Microcrystalline cellulose with an average particle size of 90 µm (Xylem Analytics, Weilheim, Germany) was used as positive control, and negative controls without bioplastic were also included. The bottles were incubated with agitation in the dark at 15°C for 28 days and then processed immediately.

### Biodegradation measurements

*In situ* biodegradation was estimated based on the weight attrition of the bioplastic sheets. The sheets were immersed in 2% sodium dodecyl sulfate and incubated overnight at 35°C with mild agitation, then brushed gently, rinsed with ultrapure water, and dried overnight at room temperature. The sheets were then weighed using a semi-micro balance (EP-225-SM-DR), and the weight attrition was calculated as the difference between the initial and post-incubation weights divided by the initial weight. Changes in the surface chemical structure of the bioplastic sheets were assessed using an Agilent 4300 Fourier transform infrared (FTIR) spectrometer (Agilent, Santa Clara, USA) in the spectral range of 650–4000 cm^−1^. The spectra were compared to reference data from the Polymers and Polymer Additives Spectra Database P/N 30002 with *MicroLab* v5.8 [38]. *In vitro* bioplastic biodegradation was estimated according to the D6691-17 standard [39] using a WTW OxiTop-IDS12 respirometer (Xylem Analytics, Weilheim, Germany), which records biological oxygen demand (BOD) by pressure difference from the absorption of CO_2_ evolved during incubation with NaOH. BOD was converted to carbon loss to CO_2_ based on carbon contents determined using a CHN analyser (ALS, Prague, Czechia).

### Metagenome and metatranscriptome sequencing

The *in situ* bioplastic sheets were scraped using a sterile scalpel and the biofilms were transferred to 2 ml Lysing Matrix E tubes (MP Biomedicals, Irvine, USA). The *in situ* seawater and the *in vitro* samples were vacuum filtered through Whatman 0.22 µm membrane filters (Cytiva, Marlborough, USA) and transferred to 2 ml cryovials. The tubes were flash-frozen in liquid nitrogen and stored at –80°C. DNA and RNA were co-extracted using a modified cetrimonium bromide (CTAB), phenol-chloroform, and bead-beating protocol [40]. After bead-beating, the extract was transferred to a clean tube and centrifuged at 16,000 × g for 5 min at 4°C. The top aqueous layer was transferred to a clean tube containing 500 µl of ≥ 99.9% stabilised chlorophorm (VWR Chemicals, Radnor, USA), and the extraction continued as further described [40]. Blank samples were extracted and sequenced in parallel. Library preparation and sequencing were done at the DNA Sequencing and Genomics Laboratory (University of Helsinki, Helsinki, Finland). The DNAprep kit (Illumina, San Diego, USA) with in-house Nextera primers was used for metagenomics and the NEBNext Ultra II Directional RNA kit (New England Biolabs, Ipswich, USA) for metatranscriptomics. The libraries were sequenced using the NovaSeq 6000 system (Illumina) and the S4 chemistry (2 × 150bp). In total, 391 metagenomes and 78 metatranscriptomes were sequenced, yielding 1.5 Tb and 508.6 Gb with a median output of 3.6 Gbp and 7.0 Gbp per sample, respectively (**Table S1**). Some PA, PE, and glass metagenomes and metatranscriptomes yielded few reads, and several metatranscriptomes could not be sequenced due to low RNA yield.

### Taxonomic profiling

*Cutadapt* v4.4 [41] was used to trim sequencing adapters, low quality base calls (Phred score < 28), and poly-A tails, and to discard short reads (< 50 bp after trimming) and reads containing ambiguous base calls. The quality of the raw and filtered data was assessed with *FastQC* v0.12.1 [42] and *MultiQC* v1.15 [43]. The filtered metagenomic and metatranscriptomic reads were classified at the domain level using *Kaiju* v1.10.0 and the *nr_euk* database release 2023-05-10 [44]. A more detailed taxonomic profiling of bacteria and archaea was obtained based on 59 single-copy marker genes with *SingleM* v0.16.0 [45] and the genome taxonomy database (GTDB) release 214 [46]. A consensus taxonomic profile was obtained by summarizing the coverage of each taxon across the 59 marker genes.

### Metagenome assembling and binning

The metagenomes were assembled and binned using the *snakemake* metagenomics workflow in *anvi’o* v7.1 [47]. For each group of 3 or 5 replicates (*in vitro* an *in situ* incubations, respectively), the filtered reads were pooled and assembled using *MEGAHIT* v1.2.9 [48] with a minimum contig length of 1000 bp, and gene calls were identified with *Prodigal* v2.6.3 [49]. Taxonomy was assigned to the contigs based on a set of 71 bacterial and 76 archaeal single-copy genes [50] identified with *HMMER* v.3.3.2 [51] and classified with *DIAMOND* v2.1.7 [52] against the GTDB release 89 [46]. The metagenomic reads were mapped to the contigs with *Bowtie2* v2.3.5 [53] and *SAMtools* v1.10 [54], and *MetaBAT2* v2.15 [55] was used to bin metagenome-assembled genomes (MAGs) based on tetranucleotide frequency and differential coverage. The MAGs from the different assemblies were pooled and pairwise average nucleotide identity (ANI) values were computed with *FastANI* v1.3.2 [56]. For each cluster of MAGs representing closely related populations with ≥ 98% ANI, a representative MAG with the highest score was selected, where

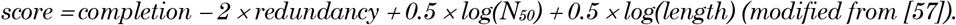

The non-redundant MAGs were visually inspected with *anvi’o* v8.0 [47] and the contigs with discrepant coverage, tetranucleotide frequency, or taxonomic signal were removed [58]. The MAGs were classified against the GTDB release 214 [46] with *GTDB-Tk* v2.3.0 [59]. The metagenomic and metatranscriptomic reads were mapped to the curated MAGs with *Bowtie2* v2.5.1 [53] and *SAMtools* v1.18 [54], and *CoverM* v0.7.0 [60] was used to count the number of reads recruited by each MAG applying the reads per kilobase per million mapped reads (RPKM) transformation. Only matching reads with ≥ 95% sequence identity and ≥ 75% coverage and MAG-level horizontal coverage/detection ≥ 25% were considered to limit the effect of non-specific mapping.

### Annotation of metabolic and plastic-degrading genes

MAG-level metabolic traits were reconstructed with *microTrait* v1.0.0 [61] and the prevalence of each trait was calculated as the proportion of MAGs containing the trait relative to all the MAGs detected in a given sample. Putative plastic-degrading genes were searched against the CAZy database [62] with *HMMER* v.3.3.2 [51] (CA; e-value < 10^–12^) and the PlasticDB database [63] with *DIAMOND* v2.1.8 [52] (PBS and PHB; ≥ 30% amino acid similarity and ≥ 70% alignment). Relatively low thresholds were applied initially to capture potentially distant homologs. For each putative plastic-degrading gene, the coverage in the corresponding *in vitro* metatranscriptomes was extracted with *anvi’o* v8.0 [47], and genes with mean horizontal coverage ≥ 25% were considered active.

### Statistical analyses

Statistical analyses were done in *R* v4.3.3 [64]. Microbial richness and beta-diversity were computed based on the read count of unique variants of the *rpsB* gene, which is a single-copy housekeeping gene encoding the universal ribosomal protein S2 of bacteria and archaea. Samples with < 325 *rpsB* reads (the median value across the dataset) were discarded in both analyses and microbial richness was estimated after subsampling the dataset. Significant differences in biodegradation and microbial richness between the materials were assessed using analysis of variance (ANOVA) followed by the Sidak test using the packages *stats* v4.3.3 [64], *emmeans* v1.10.6 [65], and *multcomp* v1.4.26 [66]. For beta-diversity analyses, the *rpsB* counts were standardised using the Hellinger transformation, pairwise Bray-Curtis dissimilarities were computed, and differences in community structure were assessed by principal coordinates analysis (PCoA) and permutational multivariate ANOVA (PERMANOVA) using the package *vegan* v2.7.1 [67]. Differentially abundant genera, traits, and MAGs were identified using linear regression with the package *Maaslin2* v1.16.0 [68]. The data were log-transformed prior to model fitting, and the *P* values were adjusted using the Benjamini-Hochberg false discovery rate (FDR). Observations with adjusted *P* < 0.05 and log2 fold change (log2FC) ≥ 3 were considered differentially abundant, the latter corresponding to a ≥ 8-fold difference between the treatments.

## Results

### *In situ* and *in vitro* bioplastic biodegradation

Biofilms were visible on the CA, PBS, and PHB sheets 12 weeks after the start of the *in situ* mesocosm experiment (**Fig. 1a**). CA and PBS, in particular, developed thick, slimy biofilms with pronounced black pigmentation. Biofilms were less developed on the other bioplastic materials (CAP, PA, and PE) and the control surface (glass). *In situ* biodegradation rates based on weight attrition differed between the bioplastic materials (ANOVA, *R*^2^ = 0.99, *P* < 0.001) (**Fig. 1b**). CA degraded the most (27.8% ± 0.2% weight attrition after 97 weeks) followed by PBS (12.7% ± 1.0%) and PHB (7.1% ± 0.2%). Weight attrition was negligible for CAP (1.3% ± 0.2%) and below detection for PA and PE. A comparable pattern was observed using FTIR spectroscopy, which revealed changes in the surface chemical structure of the CA, PBS, and PHB sheets but not CAP, PA, and PE (**Fig. 1c**).

**Figure 1.**
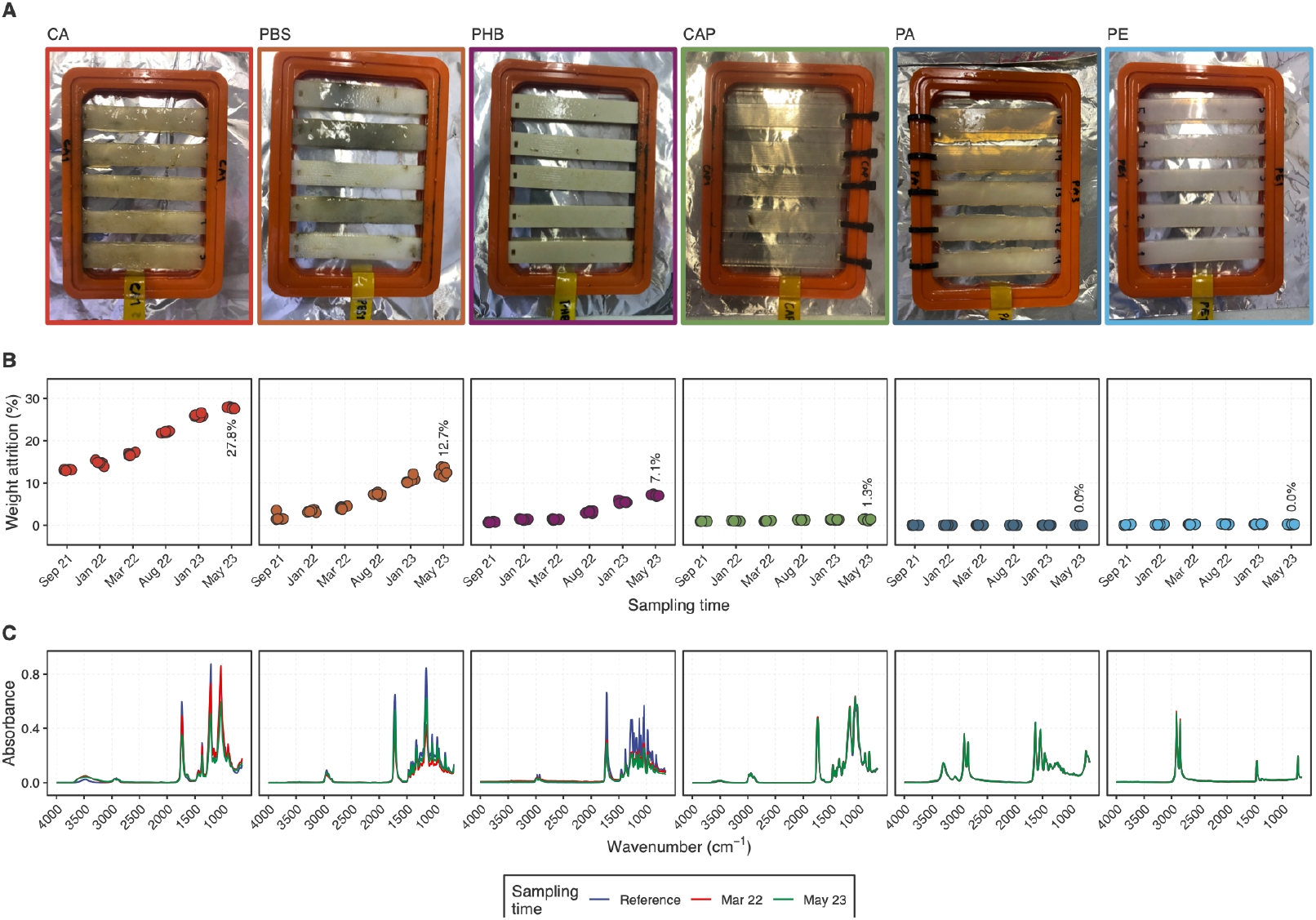
Biodegradation of six bio-based bioplastic materials in a 97-week *in situ* mesocosm experiment in the Baltic Sea. **a)** Pictures of the bioplastic sheets after 12 weeks (September 2021). **b)** Biodegradation rates based on weight attrition relative to the start of the experiment (July 2021). **c)**. Fourier transform infrared spectra of the bioplastic sheets after 38 and 97 weeks (March 2022 and May 2023, respectively).

I*n vitro* biodegradation rates based on respirometry also varied between the materials (ANOVA, *R*^2^ = 0.71, *P* < 0.001) (**Fig. S1**). PBS degraded the most (56.0% ± 7.2% carbon loss to CO_2_ after four weeks) followed by CA (32.8% ± 2.2%) and PHB (26.8% ± 11.5%). Negligible carbon loss was measured for PA and PE (1.8% ± 1.4% and 1.1% ± 1.0%, respectively), likely reflecting the utilisation of residual plastic additives. The *in vitro* biodegradation of CAP was not assessed. Temporal differences in biodegradation were observed for some materials (ANOVA, *R*^2^ = 0.95–0.99, *P* < 0.001). Whereas CA degraded uniformly, PBS varied from 7.0 ± 0.0% to 62.7% ± 4.7% and PHB from 10.7% ± 0.6% to 44.3% ± 5.0%. PBS and PHB had the highest biodegradation for communities sampled in winter (January 2022 and 2023, respectively).

### Microbial communities associated with the bioplastics

In line with our hypothesis, the analysis of reads matching the *rpsB* gene showed that the metagenomes were structured by material (PERMANOVA, *R*^2^ = 0.26, *P* < 0.001). The PCoA further clustered the metagenomes into four groups comprising *i)* the three biodegraded plastics (CA, PBS, and PHB); *ii*) the slowly degrading bioplastic (CAP); *iii*) the inert surfaces, including the bioplastics that did not degrade (PA and PE) and the control material (glass); and *iv*) the planktonic seawater metagenomes (**Fig. 2a**). The metagenomes from the biodegraded plastics and CAP had lower microbial richness than the other two groups (ANOVA, *R*^2^ = 0.63, *P* < 0.001) (**Fig. 2b**). The metagenomes were also temporally structured (PERMANOVA, *R*^2^ = 0.09–0.22, *p* < 0.001) (**Fig. S2**). The temporal dynamics varied between the biofilm and planktic datasets, with the biofilm metagenomes from both the biodegraded and non-biodegraded plastics diverging linearly over time, whereas the planktic seawater metagenomes followed a seasonal dynamic. The PHB metagenomes from January 2023 were markedly distinct from the other sampling times. Consistent microbial structure was observed for the biodegraded plastics across the *in situ* and *in vitro* metagenomes and metatranscriptomes (**Fig. S3**), which suggests that we were able to capture an active microbial community that is likely linked to the biodegradation of these materials. In contrast, the *in situ* PA and PE metagenomes clustered with the glass metagenomes, whereas the *in vitro* PA and PE metagenomes were similar to the seawater metagenomes and metatranscriptomes, suggesting activity not related to the substrate.

**Figure 2.**
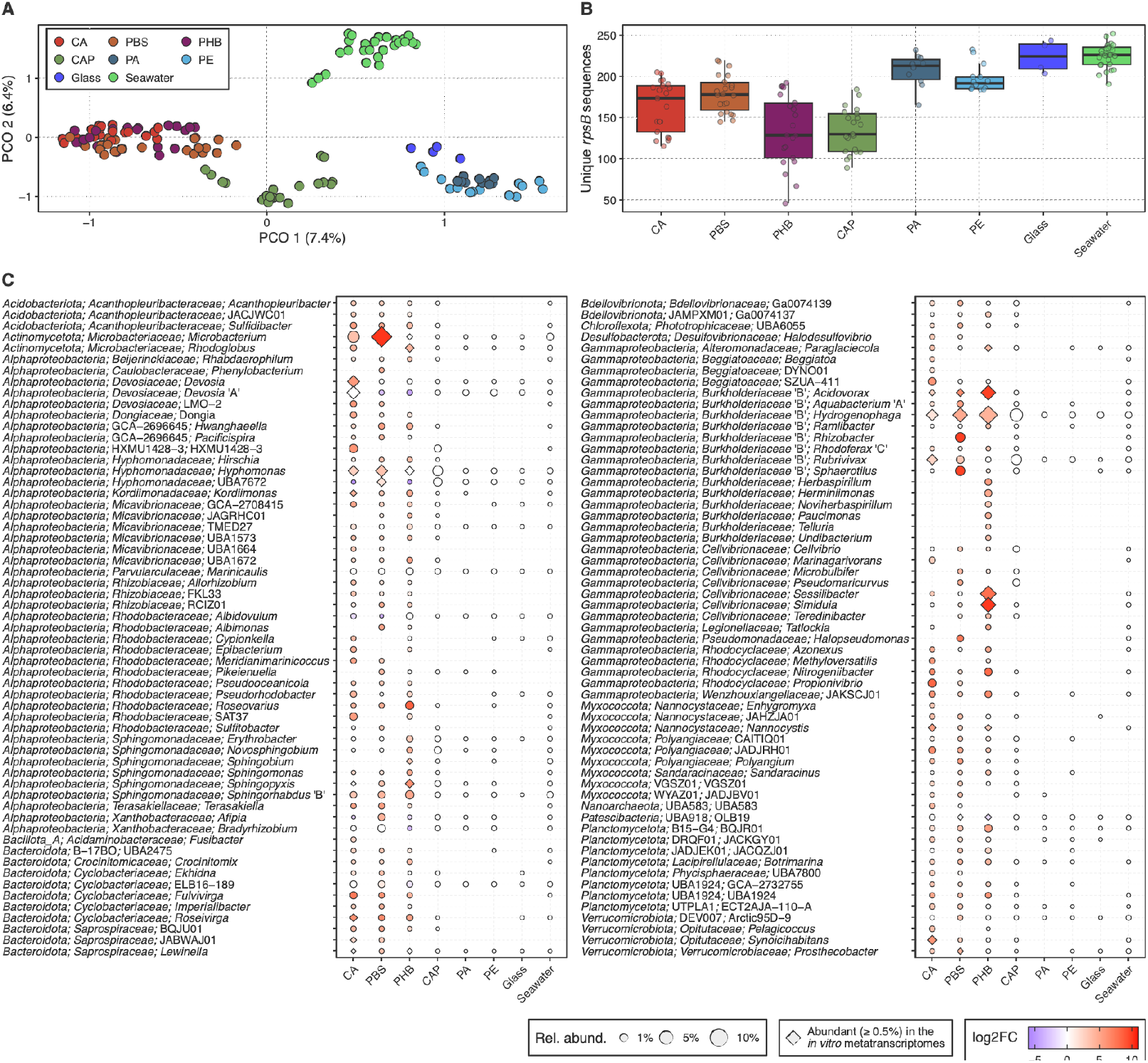
Microbial communities associated with bioplastic degradation in the Baltic Sea. **a)**. Microbial community structure based on unique 60-bp *rpsB* sequences extracted from the metagenomic reads. Principal coordinates analysis of Bray-Curtis dissimilarities of Hellinger-transformed counts.**b)**. Microbial richness based on unique 60-bp *rpsB* sequences, calculated after subsampling to even depth. In **a** and **b**, samples with < 325 *rpsB* reads were not included. **c)** Distribution of 118 abundant (≥ 0.5%) genera in the biodegraded plastics (CA, PBS, and PHB) and genera that were significantly more abundant in these materials compared to the inert surfaces (PA, PE, and glass) according to linear regression (log2FC ≥ 3.0, adjusted *P* < 0.05). Consensus taxonomic profile based on 59 marker gene sequences extracted from the metagenomic reads. Relative abundance is shown as the 10% trimmed mean across five biological replicates and six sampling times. Genera shown with a diamond were abundant (≥ 0.5%) in the associated *in vitro* metatranscriptomes.

### Potential microbial drivers of bioplastic biodegradation

Comparison with sequences from the GenBank *nr* database revealed that bacteria comprised a large fraction (98.0% ± 0.2% of the reads) of the *in situ* and *in vitro* metagenomes (**Fig. S4**). Both experiments were done in the dark to target heterotrophs and so the metagenomes contained very few reads assigned to photosynthetic microeukaryotes, which are usually abundant members of the plastisphere [30, 69]. More detailed analysis of the bacterial community based on single-copy marker genes showed that the bioplastic metagenomes were dominated by members of the *Gamma*- (26.8% ± 1.2% of the reads) and *Alphaproteobacteria* (22.6% ± 0.6%), followed by *Planctomycetota* (9.6% ± 0.4%), *Actinomycetota* (8.9% ± 0.7%), *Bacteroidota* (5.2% ± 0.2%), *Myxococcota* (2.8% ± 0.2%), *Acidobacteriota* (1.8% ± 0.1%), and *Verrucomicrobiota* (1.3% ± 0.2%) (**Fig. S5**). When considering the abundant genera—defined as those representing ≥ 0.5% of the reads in each material—only a small number of taxa were shared between the biodegraded plastic metagenomes and the other groups, and the three biodegraded plastics collectively contained the largest fraction of the total diversity (**Fig. S6**).

To explore the microorganisms that might be driving bioplastic biodegradation, we considered the abundant genera in the *in situ* CA, PBS, and PHB metagenomes and those that were significantly more abundant in these materials compared to the inert surfaces (linear regression, log2FC ≥ 3.0, adjusted *P* < 0.05). Many genera were shared between the biodegraded plastics, and many were also active in the *in vitro* metatranscriptomes (**Fig. 2c**). For example, *Hydrogenophaga* and *Acidovorax*, both members of the *Gammaproteobacteria*, and *Hyphomonas* (*Alphaproteobacteria*) were among the most abundant and active genera in the metagenomes and metatranscriptomes from the three biodegraded plastics. *Microbacterium* (*Actinomycetota*) was the most abundant genus in the *in situ* PBS and CA metagenomes and was also active in the PBS metatranscriptomes. *Devosia* (*Alphaproteobacteria*) and *Roseivirga* (*Bacteroidota*) were abundant and active in CA, and *Sessilibacter* and *Simiduia* (both *Gammaproteobacteria*) were abundant and active in PHB.

### Microbial traits associated with the bioplastics

We assembled and binned the metagenomes to investigate the functional composition of the microbial populations associated with the bioplastics. Binning yielded 5,129 non-redundant MAGs (≥ 98% ANI) which represented a substantial fraction of the metagenomes, recruiting 45.4% ± 19.4% of the reads and reaching up to 70% in some samples (**Table S1**). Metabolic reconstruction showed that the functional composition of the *in situ* metagenomes was also structured by material (PERMANOVA, *R*^2^ = 0.48, *P* < 0.001), with a clear separation between the biodegraded, non-biodegraded, and seawater metagenomes (**Fig. S7**). Differently to the results based on the *rpsB* gene, the CAP metagenomes clustered with the biodegradable plastics in the functional analysis.

We searched the MAGs for key genes involved in the initial hydrolysis of CA, PBS, and PHB to track the microbial populations that are potentially driving the biodegradation of these materials. The prevalence of bioplastic-degrading traits in the *in situ* metagenomes—defined as the proportion of MAGs containing the trait in a sample—was generally higher in the biodegraded plastics compared to the inert surfaces (linear regression, adjusted *P* < 0.05) (**Fig. 3, Fig. S8**). Genes encoding carbohydrate esterases (CAZy families CE1 and CE2) and glycoside hydrolases (CAZy families GH5, GH9, GH45, and GH48), which function in CA deacetylation and hydrolysis, respectively, were identified in 11.0% ± 0.4% of the MAGs detected in the biodegradable plastics. Genes functioning in PBS hydrolysis were identified in 11.7% ± 0.3% of the MAGs, and most were related to a PLA depolymerase derived from an uncultured microorganism with verified enzymatic activity against PBS [70]. Finally, genes encoding PHB depolymerases were identified in 40.7% ± 0.6% of the MAGs from the biodegradable plastics, 21.1% ± 2.8% from the inert surfaces and 14.3% ± 0.8 from the seawater metagenomes. Other traits that were more prevalent in the biodegraded plastics included functions related to carbohydrate depolymerization, degradation, and transport, the assimilation and transport of nitrogen, phosphorus, and sulphur compounds, aerobic respiration, anaerobic respiration, pyruvate fermentation, CO_2_ fixation, and anoxygenic photosynthesis. In contrast, lithotrophic traits, methane oxidation, and hydrogenotrophic methanogenesis were more prevalent on the inert surfaces (**Fig. 3, Fig. S8**).

**Figure 3.**
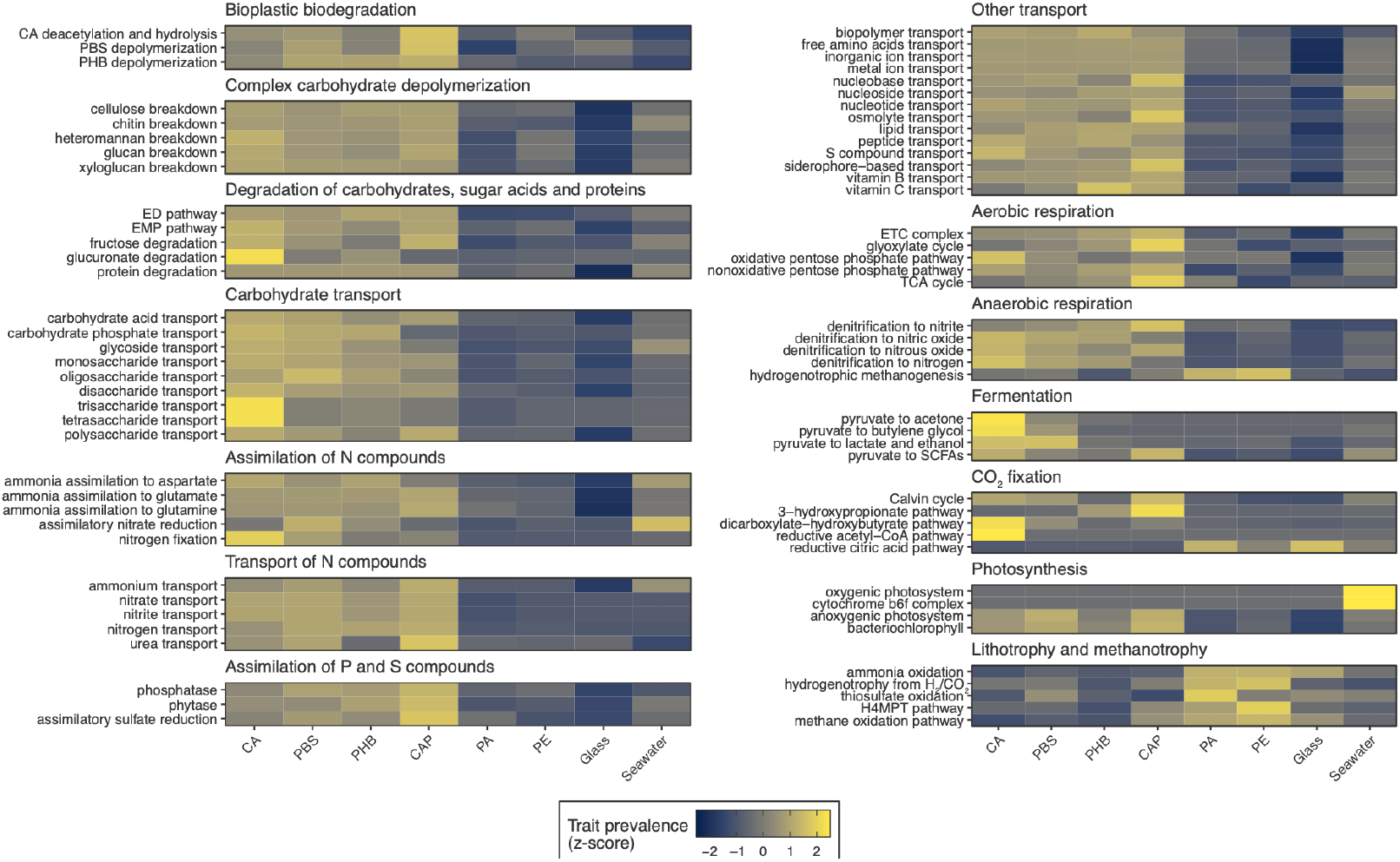
Microbial traits associated with bioplastic biodegradation in the Baltic Sea. Trait prevalence represents the proportion of metagenome-assembled genomes (MAGs) containing the trait relative to all the MAGs detected in a given sample (≥ 25% horizontal coverage). Trait prevalence is shown as the 10% trimmed mean across five biological replicates and six sampling times followed by trait-wise z-score standardisation.

### Key microbial populations in the biodegradable plastics

We investigated bioplastic biodegradation in more detail by analysing 227 medium to high quality MAGs (79.3% ± 2.1% complete, 2.8% ± 0.3% redundant) that recruited ≥ 0.5% of the reads in the CA, PBS, and PHB *in situ* and *in vitro* metagenomes and metatranscriptomes, or that were significantly more abundant in these compared to the inert materials (linear regression, log2FC ≥ 3.0, adjusted *P* < 0.05) (**Table S2**). Genes involved in the initial hydrolysis of CA, PBS, and PHB were detected in 14.5%, 18.9%, and 61.7% of the MAGs, respectively, and 19.8% encoded the genes for two or more bioplastics (**Fig. 4a**). Most MAGs with bioplastic-degrading genes belonged to *Alpha*- and *Gammaproteobacteria, Actinomycetota, Bacteroidota*, and *Myxococcota* (**Fig. 4b**).

**Figure 4.**
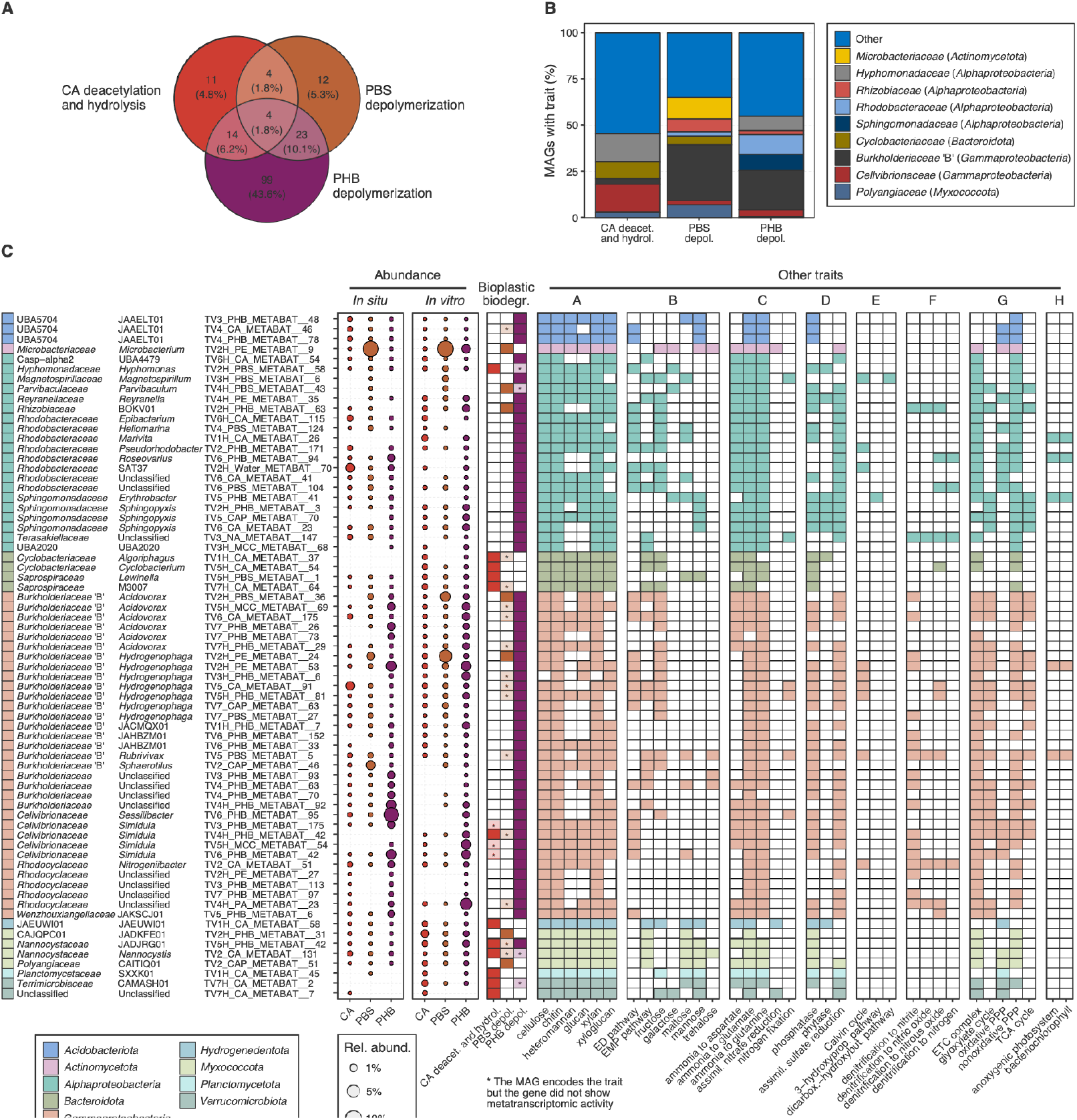
Microbial populations from the Baltic Sea with putative function in bioplastic biodegradation. Metagenome-assembled genomes (MAGs) (*n* = 227) with ≥ 0.5% relative abundance in the CA, PBS, and PHB *in situ* and *in vitro* metagenomes and metatranscriptomes, or that were significantly more abundant in these compared to the inert materials (PA, PE, and glass) according to linear regression (log2FC ≥ 3.0, adjusted *P* < 0.05). **a)** Distribution of putative plastic-degrading genes across the 227 MAGs and **b)** and their family-level taxonomic classification. **c)** Detailed information on a subset of MAGs (*n* = 69) which had plastic genes with activity in the *in vitro* metatranscriptomes. A: Complex carbohydrate depolymerization; B: Degradation of carbohydrates, sugar acids and proteins; C: Assimilation of N compounds; D: Assimilation of P and S compounds; E: CO_2_ fixation; F: Anaerobic respiration; G: Aerobic respiration; H: Photosynthesis.

The *in vitro* metatranscriptomes enabled us to probe the activity of specific genes and identify microbial populations that are likely involved in bioplastic biodegradation. Given that bioplastic was the only carbon source in the *in vitro* experiments, the metatranscriptomic signal of key bioplastic-degrading genes provides strong indication of microbial populations that are actively participating in the biodegradation of these materials. Metatranscriptomic activity was observed for 36.3%, 16.3% and 42.9% of the MAGs carrying the genes for CA, PBS, and PHB hydrolysis, respectively (**Fig. 4c, Table S2**). Many MAGs belonged to genera that were abundant in the biodegraded plastics according to the analysis of single-copy marker genes (**Fig. 2c**). Metatranscriptomic activity for PBS was detected in abundant MAGs assigned to *Microbacterium* (*Actinomycetota*), *Acidovorax*, and *Hydrogenophaga* (both *Gammaproteobacteria*), as well as in less abundant *Alphaproteobacteria* and *Myxococcota* MAGs (**Fig. 4c, Fig. 5**). In addition to PBS, the *Acidovorax* and *Hydrogenophaga* MAGs also had active genes for PHB degradation. The two *Nannocystaceae* (*Myxococcota*) MAGs and one *Simiduia* (*Gammaproteobacteria*) MAG encoded the potential to degrade the three bioplastics, but metatranscriptomic activity for PBS was not detected. Other MAGs with metatranscriptomic activity for CA included members of the *Bacteroidota* and *Verrucomicrobiota*, and MAGs with activity for PHB were diverse.

**Figure 5.**
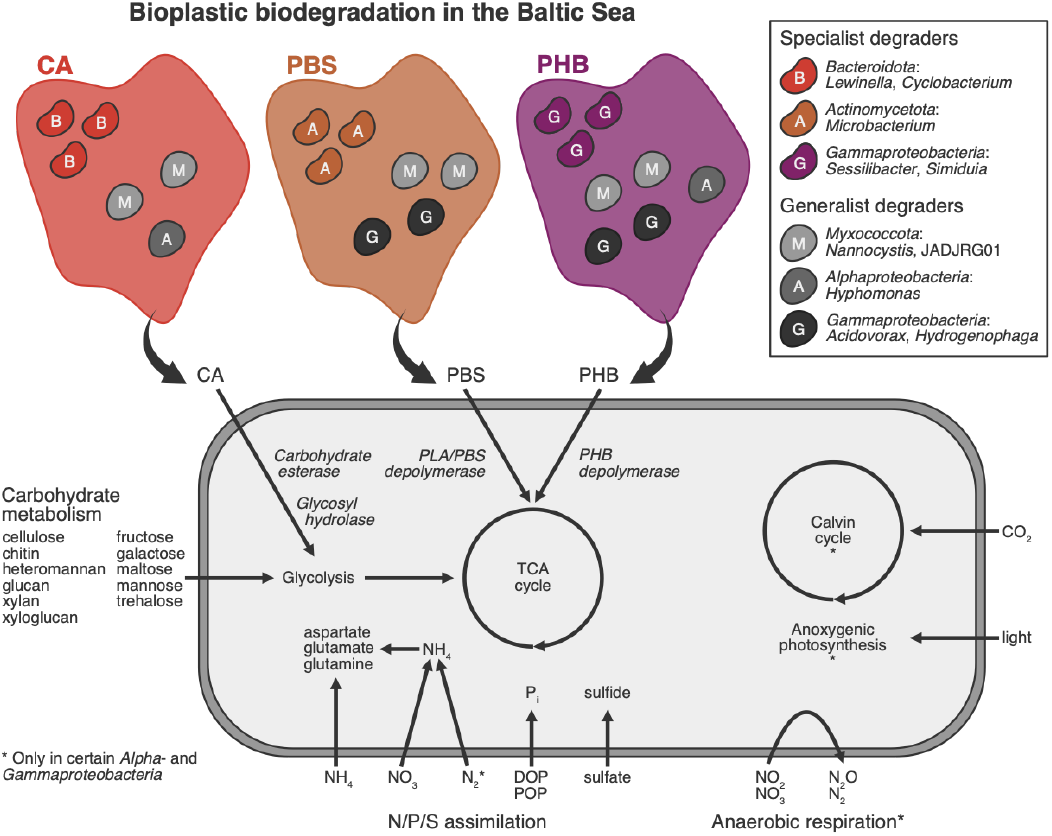
Conceptual illustration of bioplastic biodegradation in the Baltic Sea. Examples of abundant populations on CA, PBS, and PHB with metatranscriptomic activity of plastic-degrading genes, including populations of specialist and generalist degraders (active on one or many substrates, respectively). The metabolic diagram represents the aggregated metabolism of different populations with putative function on bioplastic biodegradation.

Metabolic reconstruction suggested that the populations with a putative function on CA, PBS, and PHB degradation comprised mostly aerobic heterotrophs with broad carbohydrate-degrading capabilities, including the breakdown of cellulose, chitin, heteromannan, glucan, xylan and xyloglucan (**Fig. 4c, Fig. 5, Table S2**). Several *Alpha*- and *Gammaproteobacteria* MAGs also encoded the potential for anaerobic respiration via complete and truncated denitrification (to nitrite, nitric oxide, or nitrous oxide), CO_2_ fixation (mostly via the Calvin cycle), anoxygenic photosynthesis, and nitrogen fixation.

## Discussion

Our combined experimental approach, which included multi-seasonal *in situ* and *in vitro* metagenomic and metatranscriptomic data, enabled a detailed characterisation of the biodegradation of different bioplastics in the Baltic Sea. We observed variable biodegradation rates across materials, seasons, and experimental settings. CA, PBS, and PHB degraded 7– 28% after 97 weeks *in situ* and 27–56% after four weeks *in vitro*, whereas CAP, PA, and PE had low to negligible biodegradation. In line with our hypothesis, the structure of mature (> 1 month) microbial communities differed between all the bioplastics in both taxonomic and functional spaces. However, we observed more pronounced differences between the biodegraded and non-biodedegraded plastics than within each of these groups, indicating that community development is strongly influenced by the biodegradability of the materials, as previously observed [18, 26, 71]. Community structure changed gradually from the biodegraded plastics (CA, PBS, and PHB) to the slowly-degrading plastic (CAP) and the inert materials (PA, PE, and glass), suggesting that community development on biodegradable plastics is more dynamic than on conventional plastics, in which communities are considered mature after around one month [19]. The clustering of PA and PE with glass agrees with other studies and suggests that non-biodegradable plastics act mainly as a substrate for biofilm development in marine environments, having little effect on community composition [19, 22, 26, 30, 71]. Overall, our results show that distinct biodegradable plastics can support taxonomically and functionally convergent microbial communities, which we refer to as the *bioplastisphere*.

The communities on CAP, which degraded albeit slowly, shared similarities to the communities on both the biodegraded and non-biodegraded plastics. In particular, the functional composition of the communities on CAP was similar to the bioplastisphere communities, suggesting that the biodegradation potential of CAP is comparable to the other materials, despite much slower biodegration rates. In addition to environmental drivers, the biodegradation of cellulose derivatives depends on the degree of substitution and size of the ester group [72]. The high degradation of PBS observed in our study is in contrast with previous investigations in marine and freshwater environments [25], and only the biodegradation of PBS with low molecular weight has been previsouly measured in marine environments [73]. The higher *in vitro* biodegradation rates compared to *in situ* might be attributed to experimental differences in material geometry (< 0.3 mm particles were used *in vitro* and 3 mm-thick sheets *in situ*), given that smaller plastic particles are generally easier to degrade [26, 74]. The addition of nitrogen and phosphorus in the *in vitro* incubations might have also contributed to the higher biodegradation rates. We observed variable *in vitro* biodegradation rates for each material between the different sampling times. Given that the *in vitro* experiments were done under controlled environmental settings, this suggests that the biodegradation potential of the Baltic Sea bioplastisphere fluctuates between seasons and years. Seasonal variation in biodegradation potential [26] and microbial community structure [21, 27] has been observed in previous studies. The highest biodegradation of PHB was observed in winter and coincided with a high relative abundance of *Gammaproteobacteria*, which are known to produce and degrade PHB in cold environments [75, 76]. The opportunistic behaviour of *Gammaproteobacteria* is a well-known phenomenon in decaying phytoplankton blooms and other natural pulse-like sources of dissolved organic matter [77, 78]. Altogether, our results suggest that the effects of environmental conditions, material geometry, and inoculum composition should be considered when assessing marine bioplastic biodegradation.

Our genome-resolved analysis evidenced a widespread potential for CA, PBS, and PHB degradation across the bioplastisphere independently of substrate, with many abundant populations harbouring the potential to degrade multiple bioplastics. Importantly, the metatranscriptomic activity of plastic-degrading genes provided strong indication of microbial populations that can drive bioplastic biodegradation in the Baltic Sea. Populations with active plastic-degrading were generally very abundant on the biodegraded plastics, whereas the opposite is usually observed for conventional plastics [19]. Populations with active plastic-degrading genes included common members of the marine plastisphere such as *Hyphomonas* (*Alphaproteobacteria*), *Sessilibacter*, and *Simiduia* (both *Gammaproteobacteria*) [22, 26, 28, 30], but also typically terrestrial taxa such as *Microbacterium* (*Actinomycetota*), *Acidovorax*, and *Hydrogenophaga* (both *Gammaproteobacteria*) that are generally common in coastal waters and estuaries [79–81]. Our results are therefore likely to be relevant to other coastal and estuarine systems, but differences are expected considering that biogeographical and environmental factors are known to influence the composition of the plastisphere [21, 22, 27–29].

Metabolic reconstruction revealed that the active members of the bioplastisphere are comprised mainly of aerobic and facultative anaerobic heterotrophs that encode the potential to degrade complex carbohydrates and polymers. The bioplastisphere populations were also enriched in anaerobic and nitrogen cycle traits compared to the non-biodegradable plastics. Our results indicate that nitrogen fixation might be an important strategy to counterbalance nitrogen deficiency when growing on the carbon-rich bioplastic materials [16, 24, 82]. Together with the increased capacity for anaerobic respiration via denitrification, these results also suggest that biofilm development on biodegradable plastics promotes low-oxygen microhabitats that favour facultative anaerobes [16, 24]. In addition, our results indicate that most bioplastisphere denitrifier populations have truncated pathways, as has been observed for other marine as well as terrestrial ecosystems [83, 84], and included populations with putative truncated pathways leading to the release of nitrous oxide. Overall, our study suggests that the bioplastisphere populations encoding the potential to degrade bioplastics are microbial generalists adapted to a range of oxygen, carbon, and nutrient availabilities, with diverse couplings to the overall marine carbon and nitrogen cycles.

Our study indicates that bioplastic biodegradability might be on itself a stronger driver of microbial community composition than specific physicochemical properties of the plastic materials, with the carbon derived from the breakdown of different bioplastics maintaining a common bioplastisphere. We have shown that large bioplastic fragments, equivalent to common items such as beverage bottles and disposable cuttlery, degrade in brackish seawater at rates that are much slower than those required by current industrial standards regulating bioplastic labelling. There needs to be a greater emphasis on developing more robust industrial regulations that consider more broadly the issue of mismanaged bioplastic waste that ends up in the marine environment, where they might persist for years or decades causing prolonged environmental harm. Although the search for marine biodegradable plastics and further characterisation of the bioplastisphere is an important way forward to mitigate plastic pollution, attention should be put in ensuring proper disposal of plastic waste and reducing plastic production at the source, which would also alleviate energy demands and greenhouse gas emissions.

## Author contributions

**ISP:** Formal analysis, Software, Visualization, Writing – original draft, Writing – review & editing. **EE-R:** Conceptualization, Funding acquisition, Investigation, Writing – original draft, Writing – review & editing. **PN:** Investigation, Writing – review & editing. **DNT:** Conceptualization, Funding acquisition, Investigation, Writing – review & editing. **HK:** Conceptualization, Funding acquisition, Investigation, Project administration, Writing – review & editing.

## Conflicts of interest

The authors declare no conflict of interest.

## Acknowledgements

We thank Anna Kangas, Anni Jylhä-Vuorio, and Elisa Sillförs for help with the experimental work; Murat Eren, Iva Veseli, Florian Trigodet, Tom Delmont, and Laura Cappelatti for interesting discussions and ideas; and Minna Maunula and the students of the Microbiologist’s Toolkit course (Autumn 2025, University of Helsinki) for insightful feedback on an earlier version of the manuscript. We acknowledge the DNA Sequencing and Genomics Laboratory (Institute of Biotechnology, University of Helsinki) for sequencing and the CSC– IT Centre for Finland for the computing infrastructure. No LLM/genAI tools were used in this study.

## Study funding

This study was supported by the Research Council of Finland (grant number 332174).

## Data availability

The metagenomic and metatranscriptomic data for this study have been deposited in the European Nucleotide Archive (ENA) at EMBL-EBI under the accession number PRJEB98035 (<u>ebi.ac.uk/ena/browser/view/PRJEB98035</u>).

## Supplementary material

**Pessi_et_al_2025_Suppl_Figs.docx:** Supplementary figures S1–S8.

**Pessi_et_al_2025_Suppl_Tables.xlsx:** Supplementary tables S1–S2.

## Notes

### Competing Interest Statement

The authors have declared no competing interest.

### Summary of Updates

Major changes include: - the Title has been changed to better reflect our main result and hypothesis; - the Introduction has been modified to clarify the motivation of the study and the knowledge gaps that were addressed; - several minor things have been clarified in the Materials and Methods, including justification for the choice of bioplastic materials and bioinformatic thresholds; - the Results section has been amended to emphasize the abundant and active populations in the bioplastics and the plastic-degrading genes; - two figures have been added, one of which conceptualises the taxonomic and functional dynamics of bioplastic biodegradation that were observed; and - the Discussion has been substantially rewritten to address several issues, including a better integration with our initial hypothesis.

