## Supplementary figures for "Bioplastic biodegradability shapes microbial communities in a coastal brackish environment"

### Supplementary figures S1–S8

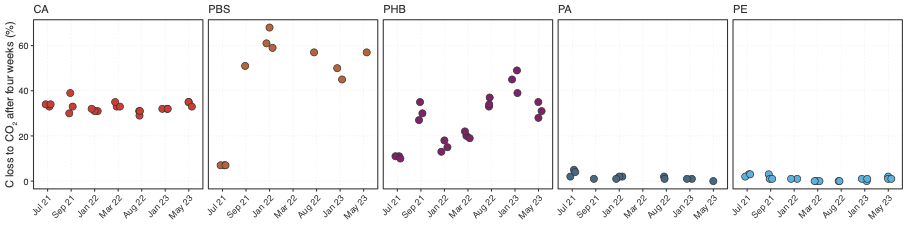

#### Figure S1

Biodegradation rates of different bioplastic materials incubated *in vitro* for four weeks with seawater from the Baltic Sea.

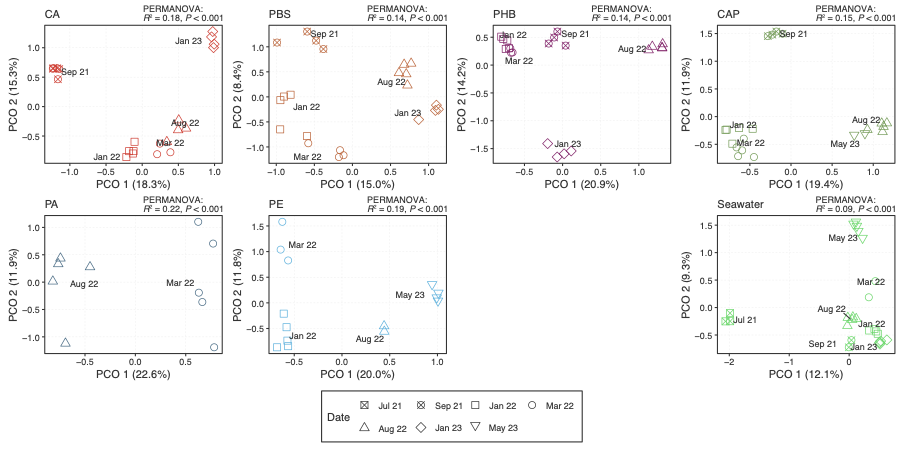

#### Figure S2

Temporal dynamics of microbial community structure in the *in situ* mesocosm. Principal coordinates analysis of Bray-Curtis dissimilarities of Hellinger-transformed counts of unique 60-bp *rpsB* sequences extracted from the metagenomic reads. Samples with
< 325 *rpsB* reads were not included.

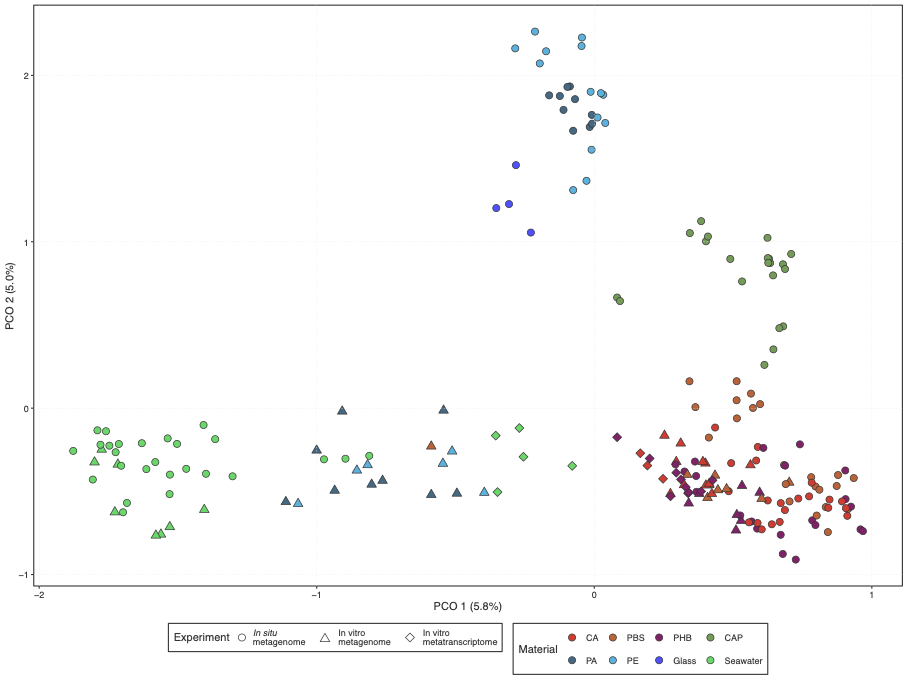

#### Figure S3

Microbial community structure across the *in situ* and *in vitro* metagenomes and metatranscriptomes. Principal coordinates analysis of Bray-Curtis dissimilarities of Hellinger-transformed counts of unique 60-bp *rpsB* sequences extracted from the reads. Samples with < 325 *rpsB* reads were not included.

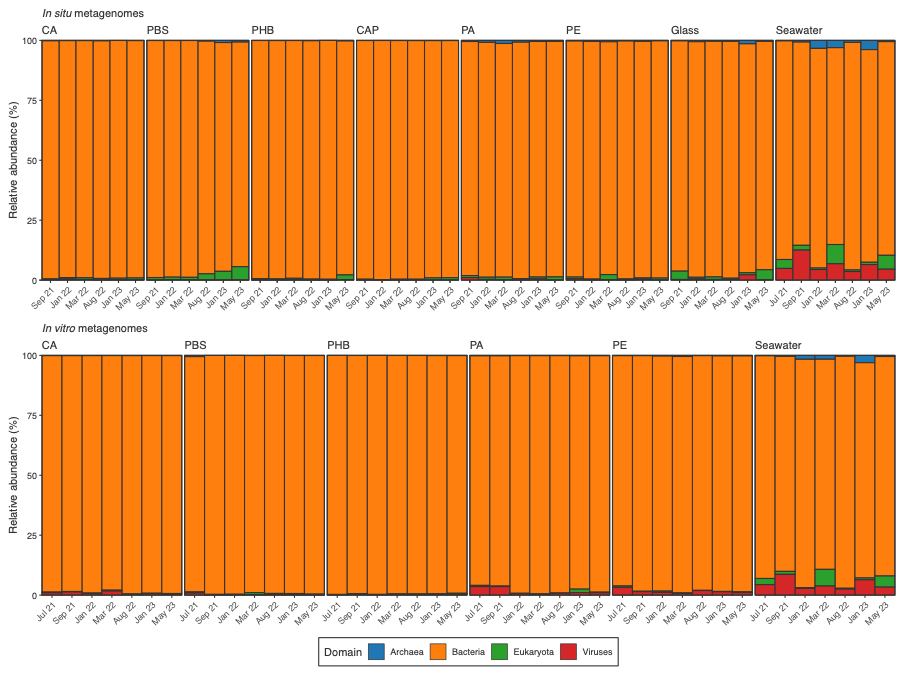

#### Figure S4

Taxonomic profiles at the domain level. Relative abundance is shown as the 10% trimmed mean across six sampling times and 3 or 5 five biological replicates (*in vitro* and *in situ* experiments, respectively).

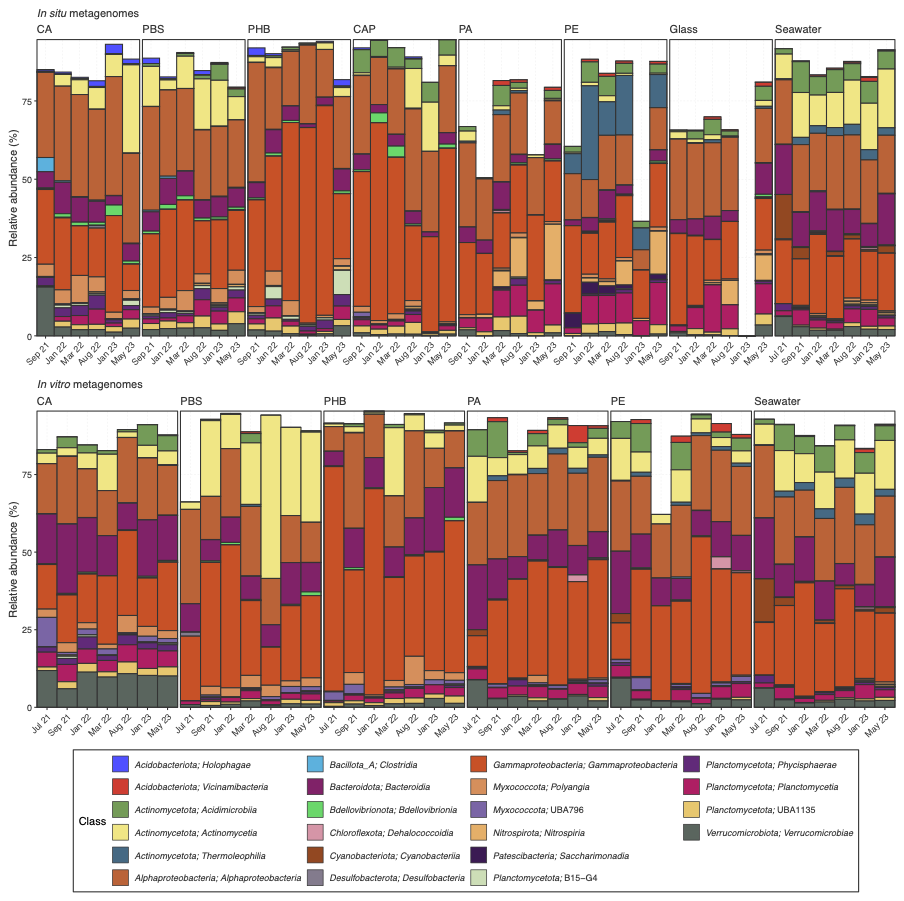

#### Figure S5

Taxonomic profiles at the class level. Consensus taxonomic profile based on 59 marker gene sequences extracted from the metagenomic reads. Relative abundance is shown as the 10% trimmed mean across six sampling times and 3 or 5 five biological replicates (*in vitro* and *in situ* experiments, respectively).

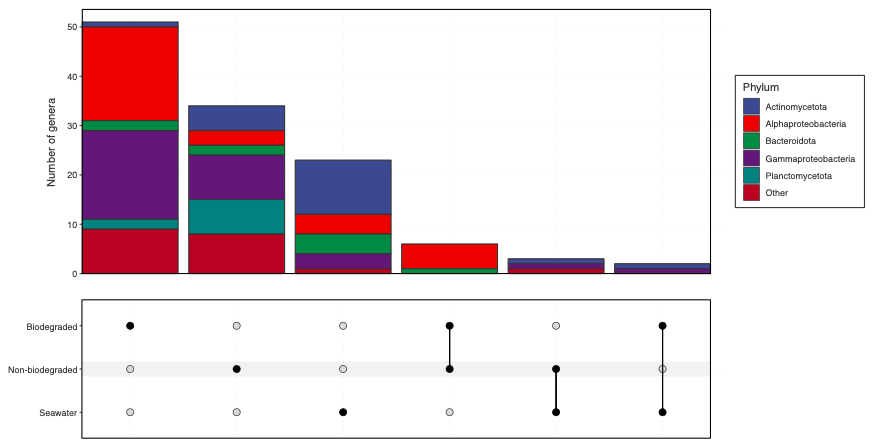

#### Figure S6

Upset plot of genus co-occurrence across the biodegraded, non-biodegraded, and seawater metagenomes from the *in situ* mesocosm. Only genera with ≥ 0.5% relative abundance in each material were included.

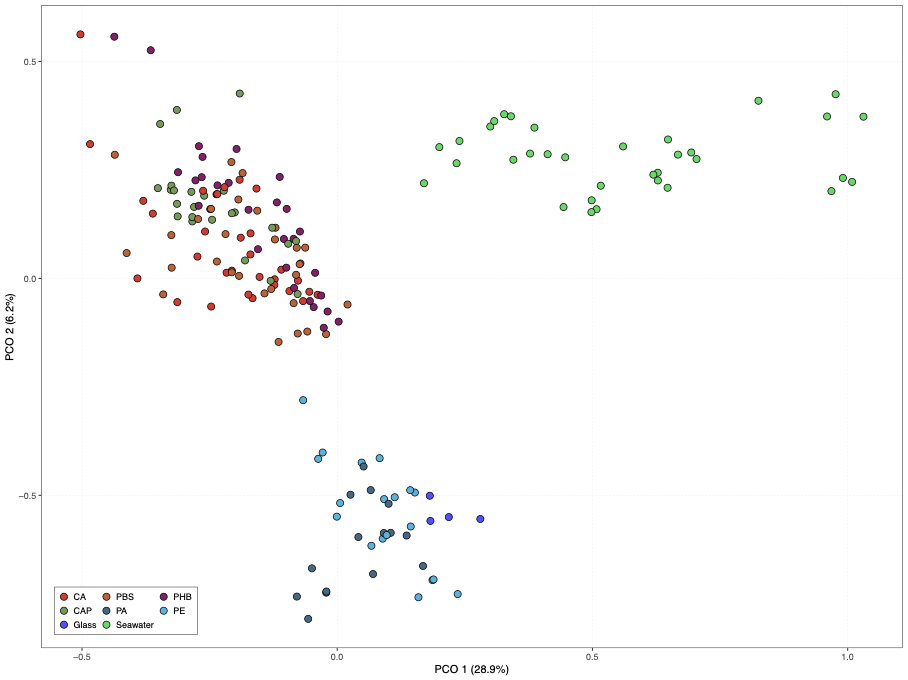

#### Figure S7

Microbial community structure in the *in situ* metagenomes in the functional space. Principal coordinates analysis of Bray-Curtis distances dissimilarities of the prevalence of metabolic traits extracted from metagenome-assembled-genomes.

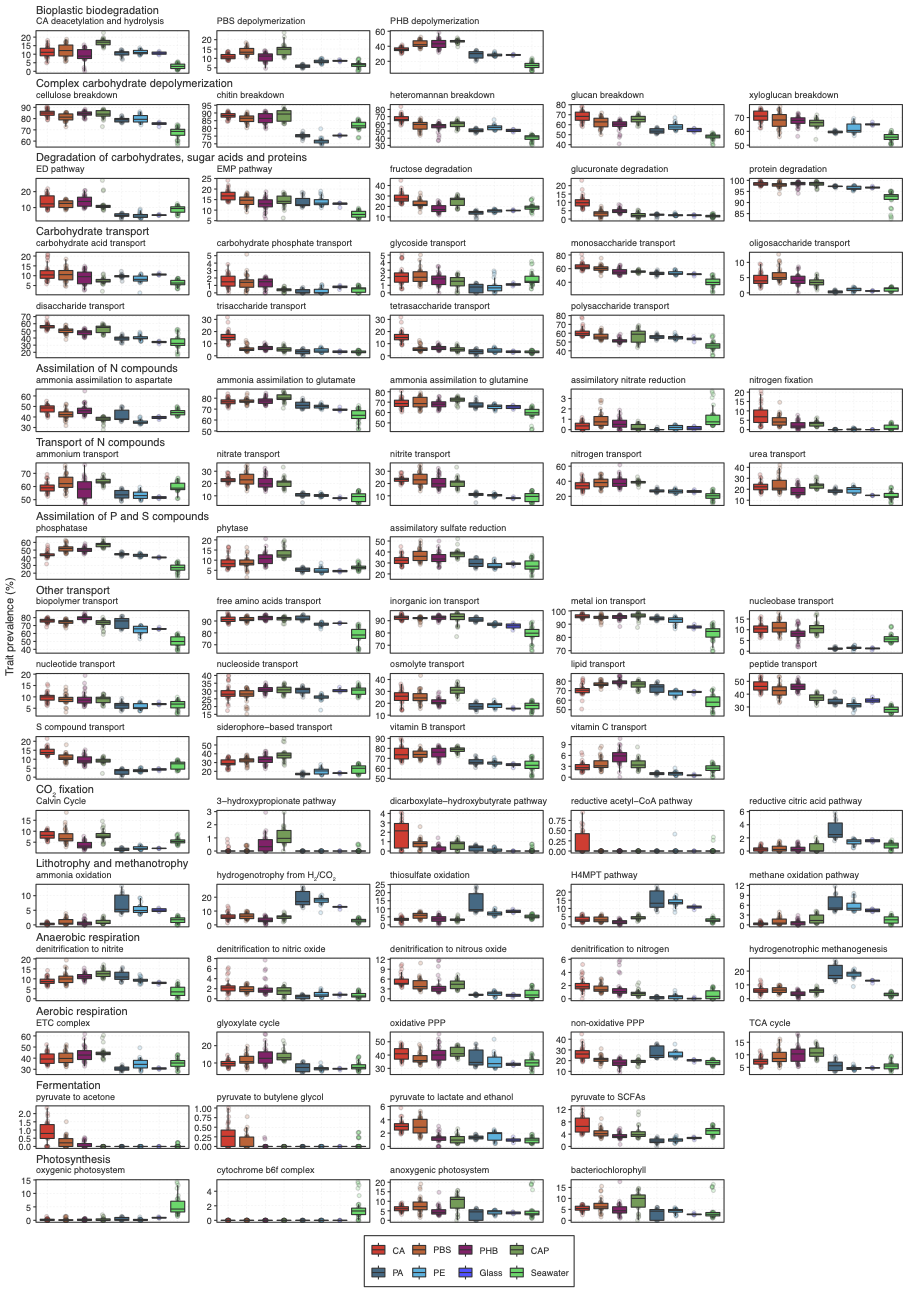

#### Figure S8

Prevalence of metabolic traits in the *in situ* metagenomes, defined as the proportion of metagenome-assembled genomes (MAGs) containing the trait relative to all the MAGs detected in a given sample (≥ 25% horizontal coverage).
